# 5-HT 2A and 5-HT 2C receptor antagonism differentially modulates reinforcement learning and cognitive flexibility: behavioral and computational evidence

**DOI:** 10.1101/2023.08.09.545287

**Authors:** Mona El-Sayed Hervig, Katharina Zühlsdorff, Sarah F. Olesen, Benjamin Phillips, Tadej Božič, Jeffrey W. Dalley, Rudolf N. Cardinal, Johan Alsiö, Trevor W. Robbins

**Author notes:** ^♱^ Joint first authors (Mona El-Sayed Hervig, Katharina Zühlsdorff). Corresponding author: Katharina Zühlsdorff Department of Psychology, Downing Place, CB2 3EB.

## Abstract

Cognitive flexibility, the ability to adapt behavior in response to a changing environment, is disrupted in several neuropsychiatric disorders, including obsessive–compulsive disorder (OCD) and major depressive disorder (MDD). Evidence suggests that flexibility, which can be operationalized using reversal learning tasks, is modulated by serotonergic transmission. However, how exactly flexible behavior and associated reinforcement learning (RL) processes are modulated by 5-HT action on specific receptors is unknown.

We investigated the effects of 5-HT_2A_ receptor (5-HT_2A_R) and 5-HT_2C_ receptor (5-HT_2C_R) antagonism on cognitive flexibility and underlying RL mechanisms. Thirty-six male Lister hooded rats were trained on a touchscreen visual discrimination and reversal task. We evaluated the effects of systemic treatments with the 5-HT_2A_R and 5-HT_2C_R antagonists M100907 and SB-242084, respectively, on reversal learning performance and performance on probe trials where correct and incorrect stimuli were presented with a third, probabilistically rewarded, stimulus. Computational models were fitted to task choice data to extract RL parameters, including a novel model designed specifically for this task.

5-HT_2A_R antagonism impaired reversal learning during certain phases. 5-HT_2C_R antagonism, on the other hand, impaired learning from positive feedback. RL models further differentiated these effects. 5-HT_2A_R antagonism decreased punishment learning rate at high and low doses. The low dose also increased exploration (beta) and increased stimulus and side stickiness (kappa). 5-HT_2C_R antagonism also increased beta, but reduced side stickiness.

These data indicate that 5-HT_2A_ and 5-HT_2C_Rs both modulate different aspects of flexibility, with 5-HT_2A_Rs modulating learning from negative feedback and 5-HT_2c_Rs for learning from positive feedback.

## INTRODUCTION

The monoamine neurotransmitter serotonin (5-hydroxytryptamine; 5-HT) system is implicated in several neuropsychiatric disorders, including major depressive disorder (MDD), obsessive– compulsive disorder (OCD) and schizophrenia, disorders in which cognitive flexibility and reinforcement learning (RL) are altered (Chamberlain et al., 2006; Clevenger et al., 2018; Zhu et al., 2021). Drugs that target the 5-HT system are often the first-line pharmacological treatment for these disorders, such as selective serotonin reuptake inhibitors (SSRIs) for MDD and OCD (APA, 2010; Fineberg et al., 2020). Emerging therapies such as the 5-HT agent psilocybin and other psychedelics are thought to hold promising treatment potential to ameliorate symptoms such as cognitive inflexibility and anhedonia (Andersen et al., 2021; Carhart-Harris & Friston, 2019; Doss et al., 2021; Stroud et al., 2018). Thus, understanding the role of serotonergic modulation mediated by specific 5-HT receptors is critical for developing future therapies for disorders characterized by inflexible behavior and diminished RL.

5-HT contributes to various cognitive processes across species, including RL (Den Ouden et al., 2013; Iigaya et al., 2018) and cognitive flexibility (Alsiö et al., 2021; Barlow et al., 2015; Clarke et al., 2004). Cognitive flexibility is defined as the ability to adapt behavior in response to changes in the environment. Inflexible behavior can manifest itself as compulsive behavior, e.g. excessively perseverative actions that are independent of outcome–value associations (Berlin & Hollander, 2014; Jentsch & Taylor, 2001; Koob & Volkow, 2016). Moreover, the ability to adjust behavior to changes in the environment is closely linked to underlying RL processes, which integrate positive and negative feedback from the environment to maximize rewards and minimize punishment (Sutton & Barto, 1998).

Flexible responding can be assessed using reversal learning paradigms across species (Uddin, 2021). During reversal learning tasks, initially learned stimulus contingencies change and the subject needs to update behavior accordingly. Substantial evidence suggests that 5-HT is involved in the modulation of reversal learning, as shown through 5-HT depletion in the orbitofrontal cortex (OFC) in monkeys (Clarke et al., 2004, 2005; Rygula et al., 2015) and rats (Alsiö et al., 2021; Izquierdo et al., 2012). In humans, acute tryptophan depletion increases outcome-independent choice perseveration (Seymour et al., 2012) and impairs reversal learning (Kanen et al., 2021). 5-HT also modulates RL processes underlying flexible behavior, possibly through distinct mechanisms (Bari et al., 2010; Seymour et al., 2012). In healthy human participants, short-term citalopram administration results in increased punishment learning and reduced reward learning (Michely et al., 2022). In patients with MDD, SSRIs impairs learning from negative feedback, while having negligible effects on learning from positive feedback (Herzallah et al., 2013). In rats, acute low-dose citalopram improves negative feedback sensitivity, while acute high-dose citalopram impairs negative feedback sensitivity, similarly to observations in human studies (Bari et al., 2010).

While it is evident that 5-HT is a key modulator of behavioral flexibility, it targets a broad range of receptor subtypes with diverse actions, exerting both excitatory and inhibitory transmission depending on receptor subtype and localization (Alvarez et al., 2021). Thus, it is vital to understand the modulatory role of 5-HT through different receptors on cognition and RL. In particular, the excitatory 5-HT_2A_Rs, which are primarily localized on excitatory pyramidal neurons, and inhibitory 5-HT_2C_Rs, found primarily on inhibitory parvalbumin neurons, seem to be involved in reversal learning – possibly with dissociable roles (Aghajanian & Marek, 1999; Amargós-Bosch et al., 2004; Liu et al., 2007; Santana et al., 2004). Systemic 5-HT_2A_R blockade impairs spatial reversal learning performance, whereas systemic blockade of 5-HT_2C_Rs improves performance (Boulougouris et al., 2008). Moreover, high levels of perseveration in rats have been found to be associated with decreased levels of 5-HT_2A_R in the OFC (Barlow et al., 2015), consistent with decreased levels of 5-HT_2A_R density in the OFC and PFC predicting clinical severity in OCD patients (Perani et al., 2008). Recent findings also suggest that psilocybin improves cognitive flexibility through a mechanism dependent on 5-HT_2A_Rs, but not 5-HT_2C_Rs (Torrado Pacheco et al., 2023). Less is known about the effects of 5-HT_2A_R and 5-HT_2C_R stimulation and blockade on component processes of reversal learning, including sensitivity to feedback and subsequent action selection.

To investigate the specific roles of 5-HT receptors in flexibility and RL, we employed the valence-probe visual discrimination (VPVD) task (Alsiö et al., 2019) and combined this task with RL modelling to gain a deeper insight into the latent processes underlying behavior. Such models are fitted to trial-by-trial data and allow for extraction of parameters such as value-dependent positive and negative learning rates, the ‘reinforcement sensitivity’ or ‘inverse temperature’ parameter, as well as the value-independent side and stimulus stickiness parameters (Daw, 2009). ‘Reinforcement sensitivity’ can also be interpreted as a measure of exploring other options (low values) versus exploiting the current likely best option given the information to date (high values). These parameters reflect different aspects of flexibility and RL, separating value-dependent from value-independent components. We examined whether these parameters contribute to choice behavior on the VPVD task and if they were affected by 5-HT_2A_R or 5-HT_2C_R blockade. We hypothesized that 5-HT_2A_R blockade would increase stickiness parameters, and that 5-HT_2C_R blockade would lead to higher learning rates, as previous studies (summarized above) have shown increased perseveration following 5-HT_2A_R blockade and improved reversal learning behavior resulting from 5-HT_2C_R antagonism. Computational modelling thus enables us to investigate the roles of the different 5-HT_2_ receptors more precisely in different aspects of RL behavior.

## MATERIALS AND METHODS

### Animals

Subjects were male hooded Lister rats (N = 36; Charles River, UK) (**Figure 1**) housed in groups of three or four throughout the experiments. The rats underwent two experiments. In the first experiment (5-HT_2A_R antagonism), all 36 rats were included. In the following 5-HT_2C_R antagonist experiment, 35 rats were included, as one rat had to be euthanized due to seizures. The rats were housed under a reverse 12-h light/dark cycle with lights off at 0700 hours. All training and testing was performed during the dark phase. To ensure sufficient motivation for task performance, the animals were food restricted with *ad libitum* access to water and fed once daily at random times after testing. Their body weights were maintained at 85% of their free-feeding weight. All experiments were subject to regulation by the United Kingdom Home Office (PPL 70/7548) in accordance with the Animals (Scientific Procedures) Act 1986.

**Figure 1.**
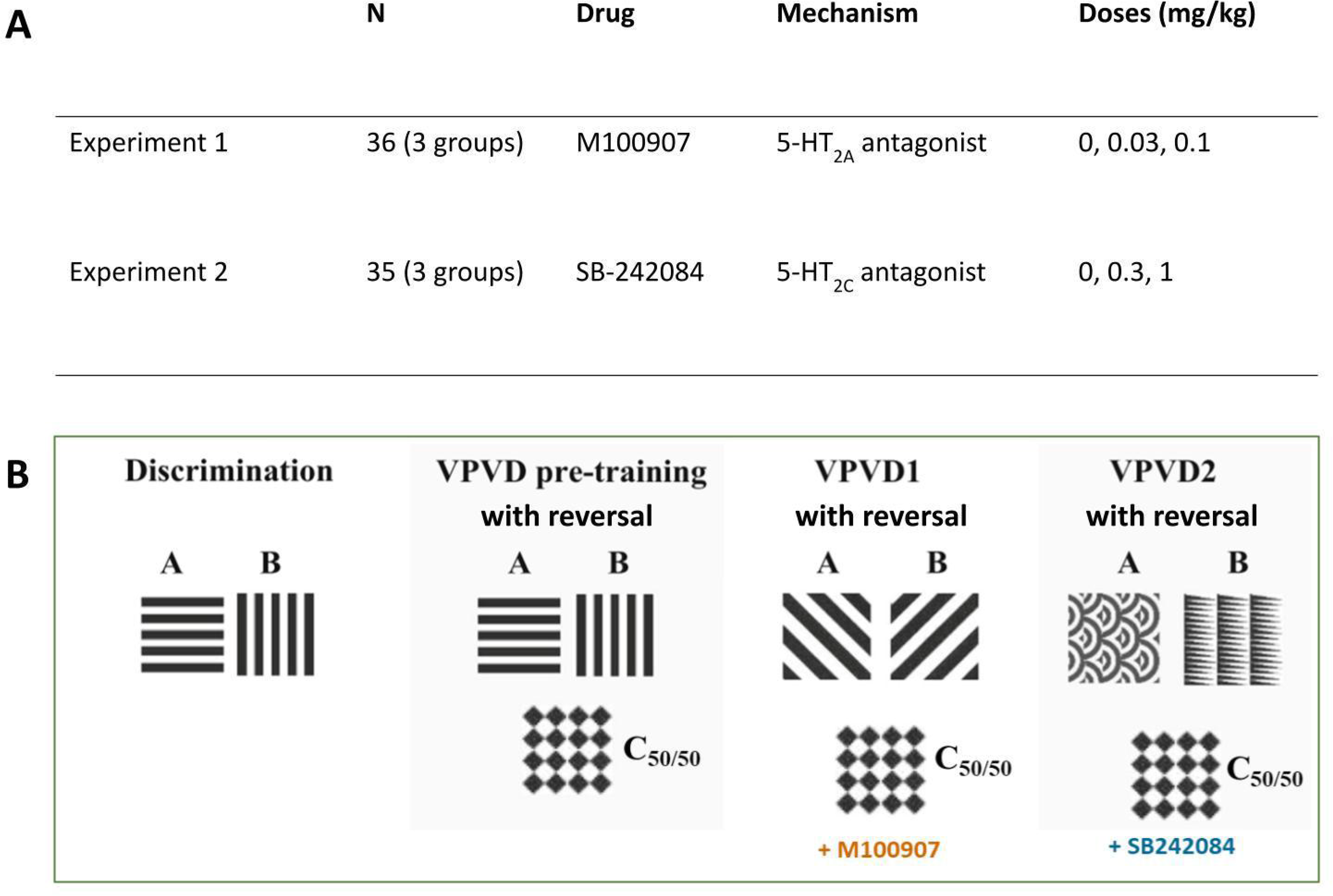
Experimental design. **A)** Table of groups and treatments. (N, number of subjects). **B)** VPVD stages and stimuli in the M100907 and SB-242084 experiments. A is the 100% reinforced stimulus, B is the 0% reinforced stimulus, C is reinforced on 50% of probe trials.

### Valence-probe visual discrimination task with reversal

Behavioral training was performed as previously described in (Alsiö et al., 2019). For experimental timeline and design see **Figure 1** and for additional information on the drugs, apparatus, behavioral pre-training, and touchscreen visual discrimination and reversal, see **Supplementary Materials**.

After pre-training, the rats progressed to the VPVD task. The VPVD task was a three-stimulus task, during which responses to one stimulus (A+) were rewarded, whereas responding to the other stimulus (B−) was punished with a time-out. A third stimulus, probabilistically rewarded on average 50% of the time (C_50/50_), was paired with either the A+ or B− on ‘probe’ trials (**Figure 1**).

The trial structure was kept constant, but a tone was played every time a trial was rewarded, and the stimulus duration was unlimited to ensure that animals completed the probe trials. The probe stimulus and frequency of probe trials (every 4 or 5 trials) were determined based on a previous study (Alsiö et al., 2019). After optimization, each of the probe trials was presented once every 8 trials: randomized, but never on the first trial within any 8-trial bin. There was a maximum of 200 trials per session. Both the inter-trial interval and time-out (on non-rewarded trials) were 5 s. Rats were initially tested for 5 days on the same A+ and B− as during the pre-training reversal (i.e., ‘horizontal bars’ vs. ‘vertical bars’). The animals then completed a visual discrimination with a novel pair of stimuli (‘slashes’ vs. ‘backslashes’; counterbalanced across rats). Training continued for a minimum of 5 sessions but could be extended to allow rats to reach 80% correct on the standard trials within the task. Next, the rats received a saline injection and were given a retention test session. The next day, rats were matched for stimulus–reward contingencies, performance on the probe trials before reversal and pre-training reversal performance, and randomly allocated to a drug group according to the experiment. The stimulus–reward contingencies were reversed on the first day of reversal and then remained the same for the duration of that experiment. The drug was administered before testing each day. The same stimulus (‘diamonds’) was used as the probe stimulus for all rats and across each of the phases. Training during the SB-242084 experiment followed the same procedure as above but rats were trained on a new pair of stimuli (‘arcs’ vs. ‘triangles’ counterbalanced across rats; the probe stimulus was kept the same) before reversal of the new stimulus−reward contingencies. In this case, the allocation into drug groups was also balanced based on previous drug exposure.

### Hierarchical Bayesian reinforcement learning modeling

The VPVD data were modeled with RL models using a hierarchical Bayesian approach. In total, nine different models were implemented in Stan (version 2.26.1), containing different combinations of parameters. The methods and models tested are described in more detail in the **Supplementary Materials**.

Q-values were updated on each trial using the following equation:

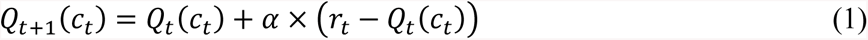

where *Q_t_*_+1_(*c_t_*) is the Q-value of the stimulus chosen on the current trial for the next, *Q_t_*(*c_t_*) is the expected value of the stimulus selected on the current trial, *α* is the learning rate and *r_t_* is the reinforcement on trial *t* (1 for reward and 0 for punishment). The learning rate reflects how much the Q-value is updated based on the prediction error *r_t_* − *Q_t_*(*c_t_*), with higher α driving faster learning.

Next, the softmax decision rule was used to calculate the probability of making one of two choices:

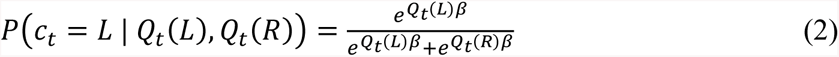

Q_t_(L) and Q_t_(R) are the Q-values of the left and right stimuli, and β is the reinforcement sensitivity parameter, which determines to what extent the subject is driven by its reinforcement history (versus random choice). Lower values of β indicate greater exploration, whereas greater values represent increased exploitation.

The behavioral data were simulated with the posterior group mean parameters from the winning model, to ensure that the model could reproduce behavioral observations. The simulations were then analyzed using a conventional approach as described below.

### Statistical analyses

Data across days within one reversal were collapsed, and trial outcomes were coded as perseverative, random, or late learning depending on performance over bins of 30 trials in a rolling window, as described in detail and illustrated previously (Hervig, Fiddian, et al., 2020), and following binomial distribution probabilities (Jones & Mishkin, 1972).

The main measures were percentage correct responses (‘% correct’) on the standard A−<B+ trials and ‘% optimal choice’ for the negative and positive probe trials across sessions. The optimal choice percentage was defined as the percentage of trials where the highest reward-probability option was chosen. Only data up to (and including) the first block of 30 trials where a rat reached criterion (24/30 correct) were analyzed.

We also analyzed response and collection latencies. Drug effects on standard parameters were analyzed using linear mixed-effects models with the lmer package in R as described previously (Phillips et al., 2018) and as recommended for such data (Wickham, 2014). The model contained two fixed factors (dose and session or dose and phase) and one random factor (subject). When relevant, further analyses were performed by conducting separate multilevel models on ‘dose’ for each session or phase. These analyses were followed by post hoc Dunnett’s corrected pairwise comparisons with the relevant vehicle condition. Significance was set at α = 0.05.

Visualization and statistical tests were performed with R, version 4.1.2 (R Core Team, 2021). Response frequencies were square-root transformed, latencies were log transformed and probabilities were arcsine transformed to ensure normality, as confirmed with a quantile–quantile plot of residuals.

## RESULTS

### Experiment 1: Effects of systemic 5-HT_2A_R blockade on reversal learning and reinforcement learning parameters

#### Effects of systemic 5-HT_2A_R blockade on reinforcement learning processes: computational modeling

After computational modeling of VPVD choice behavior, Model 9 was the best-fitting model (**Table 1**). This model included the following parameters: α_rew_ (reward learning rate), α_pun_, (punishment learning rate), β (reinforcement sensitivity), κ_stim_ (stimulus stickiness), κ_side_ (side stickiness), and the discount factor ρ. Learning from negative feedback was decreased by both low (group difference, 0 ∉ 95% HDI) and high (group difference, 0 ∉ 75% HDI) doses of M100907. There was some evidence that low, but not high, dose M100907, also decreased the reinforcement sensitivity parameter (reflecting increased exploration) (group difference, 0 ∉ 75% HDI) and increased the stimulus stickiness parameter (group difference, 0 ∉ 75% HDI). The side (location) stickiness parameter was increased in the low dose group (group difference, 0 ∉ 95% HDI) and slightly increased in the high dose group (group difference, 0 ∉ 75% HDI). The reward learning rate and discount factor were unaffected by M100907 treatment (no group differences, 0 ∈ 75% HDI) (**Figure 2 and Table 2**). The mean and standard deviation of the novel discount factor ρ for each group can be found in **Supplementary Table 2.**

**Figure 2.**
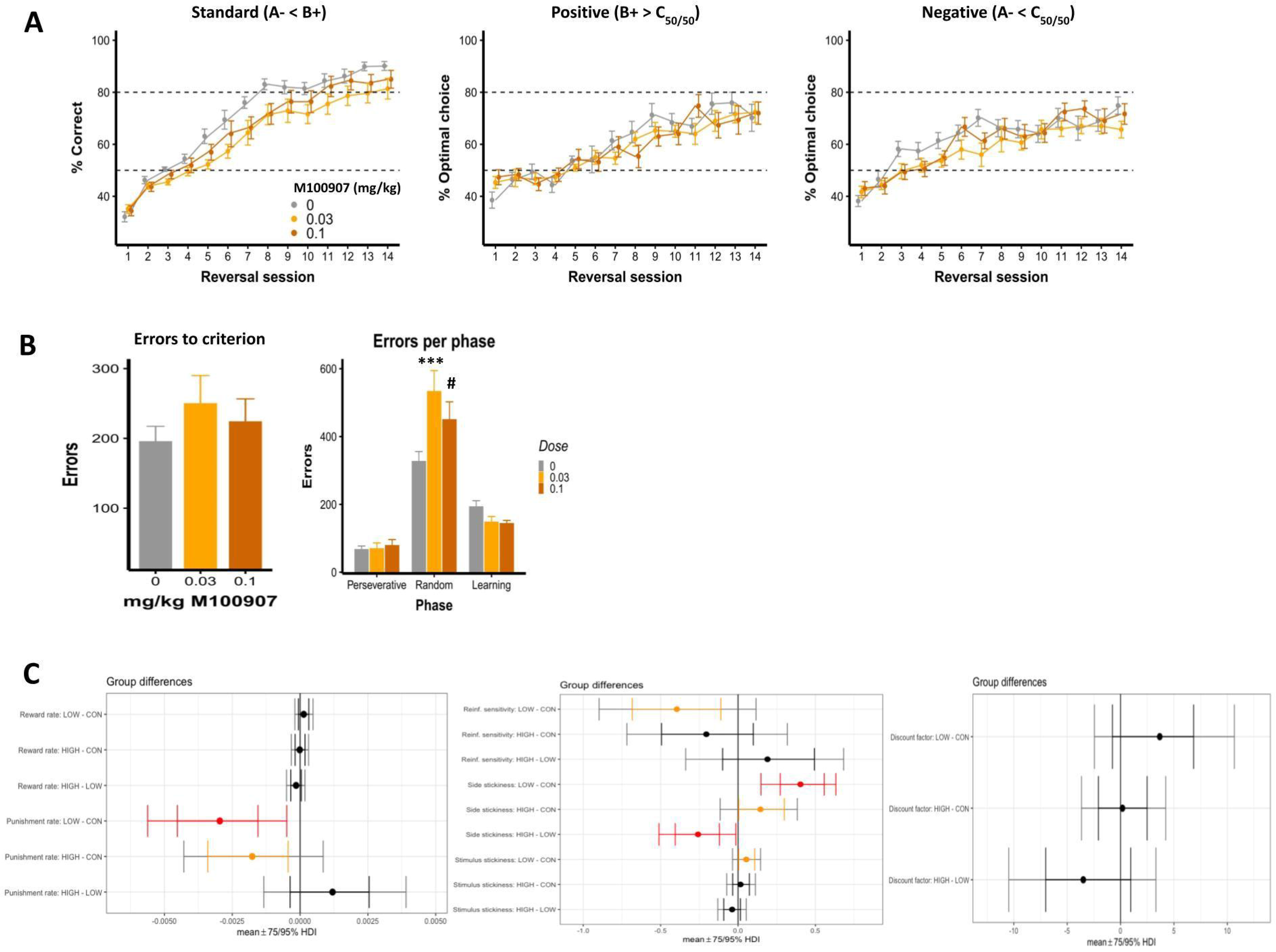
Effects of M100907 on VPVD parameters. **A)** Percent correct and percent optimal choice across sessions. **B)** Errors to criterion and errors per phase. Results are represented as mean ± standard error of the mean (SEM); *** *p* < 0.01, # p < 0.1. **C)** Results from the hierarchical Bayesian winning RL model 9, showing differences in group mean parameters following M100907 administration. (LOW, low dose; HIGH, high dose; CON, vehicle; Reinf., reinforcement; HDI, highest posterior density interval. Red indicates 0 ∉ 95% HDI; orange indicates 0 ∉ 75% HDI).

**Table 1.** Model comparison summary. Models were assumed to be equiprobable *a priori*.

| Model | Parameters | Rank (M100907) | Log marginal likelihood (M100907) | Log posterior P (M100907) | Rank (SB-248420) | Log marginal likelihood (SB-248420) | Log posterior P (SB-248420) |
| --- | --- | --- | --- | --- | --- | --- | --- |
| 1 | $\alpha, \beta$ | 9 | -57308.08 | -5267.60 | 9 | -53630.42 | -3381.41 |
| 2 | $\alpha, \beta, \kappa_{\text{stim}}$ | 8 | -57299.67 | -5259.07 | 8 | -53593.89 | -3344.88 |
| 3 | $\alpha, \beta, \kappa_{\text{side}}$ | 5 | -52320.21 | -279.61 | 4 | -50351.08 | -102.07 |
| 4 | $\alpha, \beta, \kappa_{\text{side}}, \kappa_{\text{stim}}$ | 4 | -52270.79 | -230.18 | 3 | -50322.69 | -73.68 |
| 5 | $\alpha_{\text{rew}}, \alpha_{\text{pun}}, \beta$ | 6 | -57182.04 | -5141.43 | 6 | -53561.99 | -3312.98 |
| 6 | $\alpha_{\text{rew}}, \alpha_{\text{pun}}, \beta, \kappa_{\text{stim}}$ | 7 | -57243.58 | -5202.97 | 7 | -53588.28 | -3339.27 |
| 7 | $\alpha_{\text{rew}}, \alpha_{\text{pun}}, \beta, \kappa_{\text{side}}$ | 3 | -52194.25 | -153.64 | 1 | -50249.01 | 0.000 |
| 8 | $\alpha_{\text{rew}}, \alpha_{\text{pun}}, \beta, \kappa_{\text{side}}, \kappa_{\text{stim}}$ | 2 | -52144.88 | -104.27 | 5 | -50386.28 | -127.28 |
| 9 | $\alpha_{\text{rew}}, \alpha_{\text{pun}}, \beta, \kappa_{\text{side}}, \kappa_{\text{stim}}, \rho$ | 1 | -52040.60 | 0.000 | 2 | -50255.83 | -6.82 |

**Table 2.** Summary of the effects of low and high dose M100907 and SB-242084 on reinforcement learning parameters (↑/↓, increase/decrease; - indicates no change at those levels; blank cells for parameters not tested with a given data set, as described in the Supplementary Methods). Red indicates 0 ∉ 95% HDI; orange indicates 0 ∉ 75% HDI.

| Parameter | M100907 - 0.03 mg/kg | M100907 - 0.1 mg/kg | SB-242084 – 0.3 mg/kg | SB-242084 – 1.0 mg/kg |
| --- | --- | --- | --- | --- |
| $\alpha_{rew}$ | - | - | - | - |
| $\alpha_{pun}$ | ↓ | ↓ | - | - |
| $\beta$ | ↓ | - | - | ↓ |
| $\kappa_{side}$ | ↑ | ↑ | ↓ | ↓ |
| $\kappa_{stim}$ | ↑ | - | | |
| $\rho$ | - | - | | |

Furthermore, we simulated the behavioral data using the extracted parameters from the winning model. The data modeled was separated into standard, positive and negative probe trials. The simulations were able to capture the dynamics of behavior on the VPVD task, as can be seen in the **Supplementary Materials (Figure SF.1)**.

#### Effects of 5-HT_2A_R blockade on VPVD reversal: standard behavioral parameters

There was weak evidence that systemic M100907 impaired performance on the VPVD task. On the standard (A−< B+) trials, there was a trend towards a main effect of dose (*F*_2,35_ = 2.93, *p* = 0.066) and a trend towards a dose × session interaction (*F*_26,455_ = 1.52, *p* = 0.051) (**Figure 2A**). *Post hoc* comparisons following correction for multiple comparisons revealed that the 0.03 mg/kg dose significantly reduced correct responding on sessions 6 (*t*_112_ = -2.50, *p* = 0.027), 8 (*t*_112_ = -2.63, *p* = 0.019), 13 (*t*_112_ = -2.79, *p* = 0.012) and 14 (*t*_112_ = -2.37, *p* = 0.036). On positive and negative probe trials, we found no dose × session interactions or main effect of dose.

For errors to criterion, there was a significant drug × phase interaction (*F*_4,105_ = 3.85, *p* = 0.0058), but no effect of M100907 overall (*F*_2,105_ = 0.21, *p* = 0.81). Further analysis based on planned pairwise comparisons showed that 0.03 mg/kg M100907 significantly increased errors in the mid-learning phase (*t*_115_ = 3.59, *p* = 0.0010), while there was a trend of 1 mg/kg M100907 towards increasing errors (*t*_115_ = 2.18, *p* = 0.060) in this phase.

### Experiment 2: Effects of systemic 5-HT_2C_R blockade on reversal learning and reinforcement learning parameters

#### Effects of systemic 5-HT_2C_R blockade on reinforcement learning processes: computational modeling

Model 7 was the winning model for this dataset (including parameters α_rew_, α_pun_, β and κ_side_) (Model 9 did not converge; see **Supplementary Material**). It showed that learning from positive and negative feedback were unaffected by SB-242084 (no group differences, 0 ∈ 75% HDI) (**Figure 3 and Table 2**). High-dose SB-242084 decreased the reinforcement sensitivity parameter (i.e., increasing exploration) (group difference, 0 ∉ 75% HDI). The side stickiness parameter was decreased by low-dose (group difference, 0 ∉ 95% HDI) and high-dose (group difference, 0 ∉ 75% HDI) SB-242084. We also simulated the data for this experiment using the extracted parameters (**Figure SF.2**).

**Figure 3.**
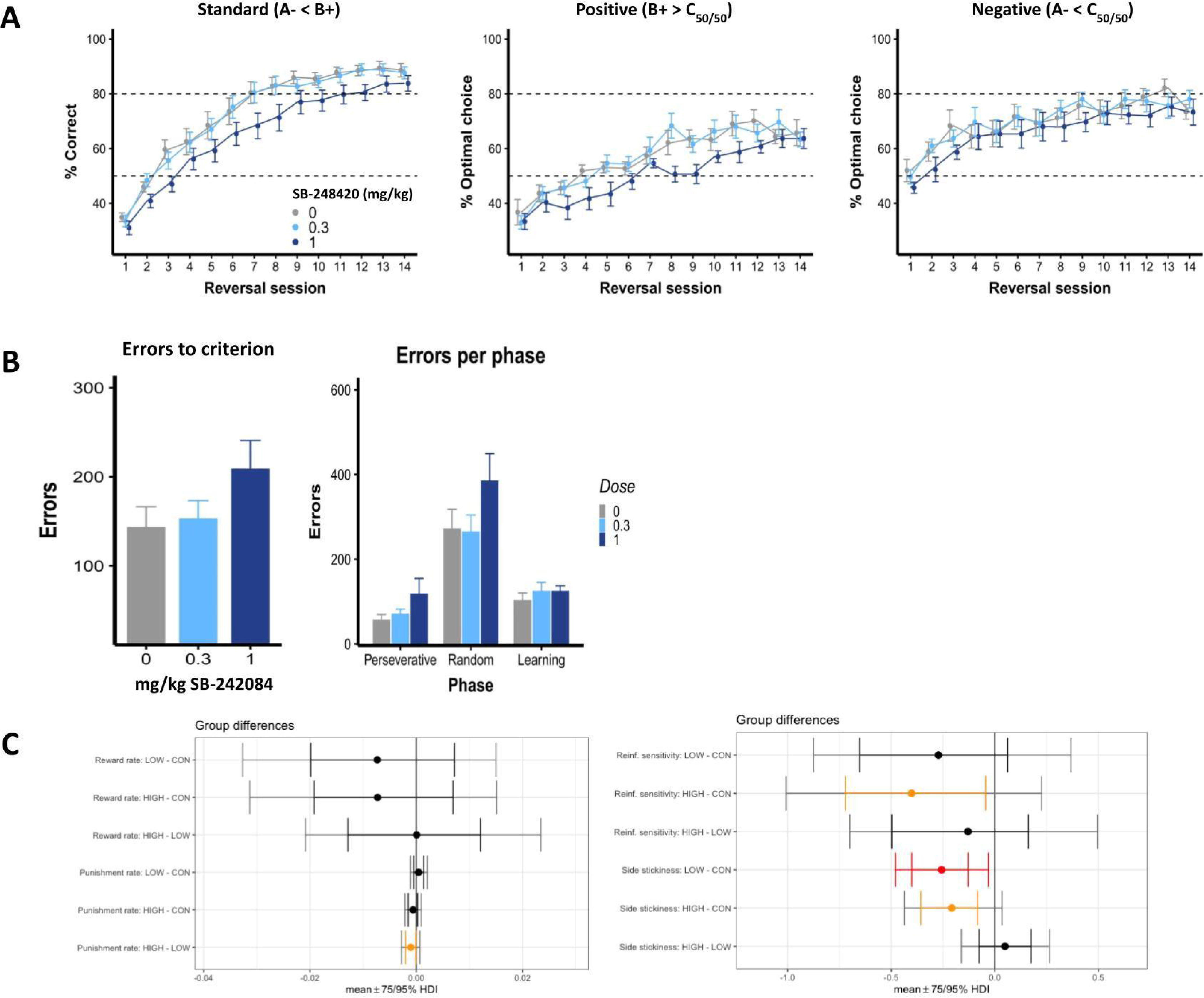
Effects of SB-242048 on VPVD parameters. **A)** Percent correct and percent optimal choice across sessions. **B)** Errors to criterion and errors per phase. Results are represented as mean ± SEM; *** *p* < 0.01, # p < 0.1. **C)** Results from the hierarchical Bayesian winning RL model 7, showing differences in group mean parameters following SB-242048 administration. (LOW, low dose; HIGH, high dose; CON, vehicle; Reinf., reinforcement; HDI, highest posterior interval. Red indicates 0 ∉ 95% HDI; orange indicates 0 ∉ 75% HDI).

#### Effects of 5-HT_2C_R blockade on VPVD reversal: standard behavioral parameters

Systemic SB-242084 impaired performance in the VPVD reversal learning task. On the standard (A−< B+) trials, there was a trend towards a main effect of dose (*F*_2,35_ = 3.15, *p* = 0.055) but no dose × session interaction (*F*_26,455_ = 0.81, *p* = 0.74) (**Figure 3**). On positive probe trials, there was a significant main effect of dose on % correct (*F*_2,35_ = 7.38, *p* = 0.0021) but no dose × session interaction (*F*_26,455_ = 1.04, *p* = 0.41).

*Post hoc* comparisons revealed that the 1.0 mg/kg SB-242084 significantly reduced % correct on sessions 7 (*t*_91.8_ = -2.63, *p* = 0.020) and 8 (*t*_91.8_ = -2.35, *p* = 0.040). On positive probe trials, % optimal choice was significantly decreased on sessions 8 (*t*_423_ = -2.48, *p* = 0.026), 9 (*t*_423_ = -2.61, *p* = 0.018), 11 (*t*_423_ = -2.39, *p* = 0.034) and 12 (*t*_423_ = -2.24, *p* = 0.049).

For errors to criterion, we found no effect of SB-242084 overall (*F*_2,105_ = 1.80, *p* = 0.17). When analyzing effect of SB-242084 on errors per phase, we found a trend towards a main effect of dose (*F*_2,35_ = 3.15, *p* = 0.055) and significant effect of phase (*F*_2,70_ = 53.15, *p* < 0.0001), but no dose × phase interaction (*F*_4,70_ = 0.50, *p* = 0.73).

Win-stay/lose-shift and latency analyses for both experiments can be found in the **Supplementary Materials**.

## DISCUSSION

These findings indicated contrasting, as well as common, effects of 5-HT_2A/C_R antagonists on measures of RL and cognitive flexibility in the rat. We used a computational modeling approach to visual discrimination reversal that proved more sensitive to characterizing drug effects than standard behavioral measures. The RL parameters enabled us to gain a deeper insight into the latent mechanisms underlying behavior on the VPVD task.

### Effects of 5-HT_2A_R antagonism on reinforcement learning and cognitive flexibility

Selective blockade of 5-HT_2A_Rs using M100907 impaired reversal learning as reflected by reductions in % correct on standard trials and an increasing frequency of errors at certain learning phases. This impairment was not associated with changes in response or collection latencies, showing that it was unlikely to be caused by motivational or sensorimotor deficits. Computational analyses revealed that 5-HT_2A_R antagonism impaired learning from negative feedback, decreased the reinforcement sensitivity parameter (i.e., increased ’exploration’) and increased both side and stimulus ’stickiness’, suggesting differential effects of 5-HT_2A_R blockade on value-dependent (reinforcement sensitivity) compared to value-independent (stickiness) choices, which may hence reflect distinct facets of the cognitive flexibility construct.

Previous studies using systemic (Boulougouris et al., 2008) or intra-lateral OFC (Hervig, Piilgaard, et al., 2020) M100907 have also shown impaired reversal learning performance, consistent with the present findings. Moreover, lower 5-HT_2A_R binding in the rat OFC is associated with more perseveration during spatial reversal (Barlow et al., 2015). Our findings may seem inconsistent with studies showing that the 5-HT_2A_R antagonist ketanserin normalizes impairments in flexibility resulting from lysergic acid diethylamide (LSD), as well as general improvements in set-shifting following ketanserin administration in rats (Baker et al., 2011; Pokorny et al., 2020; Torrado Pacheco et al., 2023). However, such apparent inconsistencies may have resulted from the use of different paradigms to assess flexibility, such as set-shifting, which may involve distinct neural and 5-HT dependent substrates than reversal learning (Clarke et al., 2005; Dias et al., 1996).

Dose may also be a relevant factor to consider. In our study, the lower dose of 0.03 mg/kg M100907 affected reversal learning more than the 0.1 mg/kg dose, likely reflecting an inverted U-curve effect, as previously reported for 5-HT_2A_R antagonists (Marek et al., 2005; Roth, 2011). The high 5-HT_2A_R antagonist dose may have induced receptor internalization, a known mechanism for the 5-HT_2A_R (Roth, 2011). Dose-response studies have shown that moderate systemic doses of M100907 are more effective than low and high doses on a response-inhibition task and that intra-lateral OFC infusions with moderate M100907 doses induce the most detrimental effects on reversal learning (Furr et al., 2012; Marek et al., 2005).

Our findings align with our initial hypothesis of increased stickiness following 5-HT_2A_R blockade. Selective depletions of 5-HT in the marmoset OFC and amygdala using 5,7-DHT results in increased side stickiness rates, similar to our findings, following 5-HT_2A_R antagonism (Rygula et al., 2015), suggesting that 5-HT_2A_Rs in these areas may modulate the stickiness parameter, i.e., repeating responses regardless of previous outcomes. This accords with the demonstration that side stickiness is correlated with functional connectivity between the amygdala and medial OFC in rats (Zühlsdorff et al., 2023).

### Effects of 5-HT_2C_R antagonism on reinforcement learning and cognitive flexibility

Antagonism of 5-HT_2C_Rs with SB-242084 impaired overall reversal learning performance (decreased % correct), which seemed to be driven by impaired learning from positive feedback (decreased % optimal choice on positive probe trials) on the VPVD task at high doses. This is in line with previous studies showing that 5-HT_2C_R blockade impairs late phases of visual reversal learning (Alsiö et al., 2015) and 5-HT_2C_R agonism improves flexibility (Del’Guidice et al., 2013; Phillips et al., 2018).

Using RL models, we found that 5-HT_2C_R blockade decreased the reinforcement sensitivity parameter at a higher dose and decreased side stickiness both at low and high doses. This drug therefore increased flexible responding according to different computational RL parameters. This observation is in accordance with studies showing SB-242084 to improve spatial reversal learning and early phases of serial visual reversal learning (Alsiö et al., 2015; Boulougouris et al., 2008). On a probabilistic reversal task, 1 mg/kg SB-242084 increases exploration following positive feedback, consistent with our finding of reduced reinforcement sensitivity at this dose (Phillips et al., 2018). Our findings highlight that the role of 5-HT_2C_Rs in cognitive flexibility is complex, indicating that 5-HT_2C_Rs are crucial for integrating positive feedback to optimize learning, while blocking this receptor improves flexibility as shown computationally. Thus 5-HT_2C_Rs exert different roles in reward-dependent parameters (positive feedback) versus reward-independent parameters (stickiness).

### Implications for mechanisms of action of SSRIs and psychedelics in psychiatric disorders

In a recent analysis, lower doses of the SSRI citalopram increases the reward learning rate and decreases side stickiness, whilst decreasing reward rate and increasing reinforcement sensitivity at a higher dose (Luo et al., 2023). Acute escitalopram in healthy human participants reduces the reward learning rate, decreases reinforcement sensitivity, and decreases stimulus stickiness (Luo et al., 2023), partially aligning with our findings following 5-HT_2C_R blockade. Our findings using selective 5-HT_2A_R and 5-HT_2C_R antagonists may thus aid our understanding of mechanisms underlying cognitive flexibility and RL.

Psilocybin and other psychedelics are receiving increased attention for their therapeutic potential in treating neuropsychiatric disorders such as MDD and anxiety (Carhart-Harris et al., 2016, 2021; Goldberg et al., 2020). Even though their mechanisms are poorly understood, one hypothesis is that psilocybin improves cognitive flexibility (Baker et al., 2011; Torrado Pacheco et al., 2023). Psilocybin, which primarily exerts its psychoactive effects through 5-HT_2A_R agonism (Madsen et al., 2019), has been shown to increase cognitive flexibility in individuals with MDD for at least 4 weeks (Doss et al., 2021). Ayahuasca, which contains the 5-HT_2A_R agonist dimethyltryptamine, similarly increases cognitive flexibility in healthy volunteers (Kuypers et al., 2016; Murphy-Beiner & Soar, 2020). In contrast, 2,5-dimethoxy-4-iodoamphetamine, a 5-HT2_A/C_R agonist, impairs flexible strategy choice, highlighting different mechanisms of actions of hallucinogenic substances (Torrado Pacheco et al., 2023). Finally, a recent study investigating the effects on RL parameters of the psychedelic LSD, a partial 5-HT_2A_R agonist, has reported increased reward and punishment learning, and reduced stimulus stickiness (Kanen et al., 2022). Overall, these results suggest that 5-HT_2A_R agonism can improve flexibility. In the present study, we show that antagonism of this receptor decreases the punishment learning rate and increases stickiness, mirroring these hypothetical effects of 5-HT_2A_R agonism. A limitation of our study is the fact that only male animals were included; therefore, sex-dependent effects could not be investigated.

In summary, we report that both 5-HT_2A_R and 5-HT_2C_R antagonism altered performance on a visual reversal task. We characterized this impairment using RL models, finding that 5-HT_2A_R blockade reduced both learning from punishment and reinforcement sensitivity, but increased stickiness. 5-HT_2C_R blockade impaired learning from positive feedback as assessed using conventional measures, suggesting a dissociation between the two receptors: the 5-HT_2C_R is essential for learning from positive feedback and the 5-HT_2A_R is important for learning from negative feedback. Additionally, 5-HT_2C_R antagonism reduced reinforcement sensitivity and side stickiness parameters, indicating increased flexibility. These results provide novel insights into the mechanisms of 5-HT and the involvement of different 5-HT receptors in cognitive flexibility. This may be important for our understanding of neuropsychiatric conditions such as MDD and OCD, as well as for research into future treatments such as psychedelic agents that act as 5-HT_2A_R agonists.

## Supporting information

Supplementary materials and results

## AUTHOR CONTRIBUTIONS

MEH: conceptualization, methodology, investigation, data curation, formal analysis, writing – original draft, writing – review & editing; KZ: conceptualization, software, methodology, formal analysis, writing – original draft, writing – review & editing; SFO – methodology, investigation, data curation; BP – methodology, investigation, data curation, formal analysis; TB – methodology, investigation, data curation; RNC – software, methodology, formal analysis, writing – review & editing; JWD – conceptualization, writing – review & editing; JA – conceptualization, methodology, software, formal analysis, writing – review & editing; TWR – conceptualization, methodology, writing – original draft, writing – review & editing, supervision, funding acquisition.

## FUNDING

This work was supported by a Wellcome Trust Senior Investigator Grant to TWR (104631/Z/14/Z) and a Lundbeck Foundation Research Fellowship to MEH (R182-2014-2810 and R210-2015-2982). KZ was supported by the Institute for Neuroscience at the University of Cambridge, the Alan Turing Institute, London and the Angharad Dodds John Bursary in Mental Health and Neuropsychiatry, Downing College, Cambridge. JWD has received funding from GlaxoSmithKline and Boehringer Ingelheim Pharma GmbH and is a co-investigator on an MRC program grant (MR/N02530X/1). TWR is also a co-investigator of the latter grant. RNC’s research is supported by the UK Medical Research Council (MRC) (MR/W014386/1). JA was supported by a short-term grant from Fudan University. SFO, TB and BP have no funding to declare.

## DECLARATION OF INTERESTS

JWD has received research grants from Boehringer Ingelheim Pharma GmbH and GlaxoSmithKline and receives royalties from Springer Verlag. TWR discloses consultancy with Cambridge Cognition; he receives editorial honoraria from Springer-Nature and Elsevier and a research grant from Shionogi. RNC consults for Campden Instruments and receives royalties from Cambridge Enterprise, Routledge, and Cambridge University Press. KZ, MEH, JA, SFO, TB and BP have no conflicts to declare.

## BIBLIOGRAPHY

Aghajanian, G. K., & Marek, G. J. (1999). Serotonin and hallucinogens. Neuropsychopharmacology, 21. https://doi.org/10.1016/S0893-133X(98)00135-3

Alsiö, J., Lehmann, O., Mckenzie, C., Theobald, D. E., Searle, L., Xia, J., Dalley, J. W., & Robbins, T. W. (2021). Serotonergic Innervations of the Orbitofrontal and Medial-prefrontal Cortices are Differentially Involved in Visual Discrimination and Reversal Learning in Rats. Cerebral Cortex (New York, NY), 31(2), 1090. https://doi.org/10.1093/CERCOR/BHAA277

Alsiö, J., Nilsson, S. R. O., Gastambide, F., Wang, R. A. H., Dam, S. A., Mar, A. C., Tricklebank, M., & Robbins, T. W. (2015). The role of 5-HT2C receptors in touchscreen visual reversal learning in the rat: A cross-site study. Psychopharmacology. https://doi.org/10.1007/s00213-015-3963-5

Alsiö, J., Phillips, B. U., Sala-Bayo, J., Nilsson, S. R. O., Calafat-Pla, T. C., Rizwand, A., Plumbridge, J. M., López-Cruz, L., Dalley, J. W., Cardinal, R. N., Mar, A. C., & Robbins, T. W. (2019). Dopamine D2-like receptor stimulation blocks negative feedback in visual and spatial reversal learning in the rat: behavioural and computational evidence. Psychopharmacology, 236(8), 2307–2323. https://doi.org/10.1007/s00213-019-05296-y

Alvarez, B. D., Morales, C. A., & Amodeo, D. A. (2021). Impact of specific serotonin receptor modulation on behavioral flexibility. In Pharmacology Biochemistry and Behavior (Vol. 209). https://doi.org/10.1016/j.pbb.2021.173243

Amargós-Bosch, M., Bortolozzi, A., Puig, M. V., Serrats, J., Adell, A., Celada, P., Toth, M., Mengod, G., & Artigas, F. (2004). Co-expression and In Vivo Interaction of Serotonin1A and Serotonin2A Receptors in Pyramidal Neurons of Pre-frontal Cortex. Cerebral Cortex, 14(3). https://doi.org/10.1093/cercor/bhg128

Andersen, K. A. A., Carhart-Harris, R., Nutt, D. J., & Erritzoe, D. (2021). Therapeutic effects of classic serotonergic psychedelics: A systematic review of modern-era clinical studies. In Acta Psychiatrica Scandinavica (Vol. 143, Issue 2). https://doi.org/10.1111/acps.13249

APA. (2010). Practice guideline for the treatment of patients with major depressive disorder (third edition). American Psychiatric Association.

Baker, P. M., Thompson, J. L., Sweeney, J. A., & Ragozzino, M. E. (2011). Differential Effects of 5-HT2A and 5-HT2C Receptor Blockade on Strategy-Switching. Behavioural Brain Research, 219(1), 123. https://doi.org/10.1016/J.BBR.2010.12.031

Bari, A., Theobald, D. E., Caprioli, D., Mar, A. C., Aidoo-Micah, A., Dalley, J. W., & Robbins, T. W. (2010). Serotonin modulates sensitivity to reward and negative feedback in a probabilistic reversal learning task in rats. Neuropsychopharmacology, 35(6), 1290–1301. https://doi.org/10.1038/npp.2009.233

Barlow, R. L., Alsiö, J., Jupp, B., Rabinovich, R., Shrestha, S., Roberts, A. C., Robbins, T. W., & Dalley, J. W. (2015). Markers of serotonergic function in the orbitofrontal cortex and dorsal raphé nucleus predict individual variation in spatial-discrimination serial reversal learning. Neuropsychopharmacology : Official Publication of the American College of Neuropsychopharmacology, 40(7), 1619–1630. https://doi.org/10.1038/NPP.2014.335

Berlin, G. S., & Hollander, E. (2014). Compulsivity, impulsivity, and the DSM-5 process. CNS Spectrums, 19(1), 62–68. https://doi.org/10.1017/S1092852913000722

Boulougouris, V., Glennon, J. C., & Robbins, T. W. (2008). Dissociable effects of selective 5-HT2A and 5-HT2C receptor antagonists on serial spatial reversal learning in rats. Neuropsychopharmacology, 33(8). https://doi.org/10.1038/sj.npp.1301584

Carhart-Harris, R., Bolstridge, M., Rucker, J., Day, C. M. J., Erritzoe, D., Kaelen, M., Bloomfield, M., Rickard, J. A., Forbes, B., Feilding, A., Taylor, D., Pilling, S., Curran, V. H., & Nutt, D. J. (2016). Psilocybin with psychological support for treatment-resistant depression: an open-label feasibility study. The Lancet Psychiatry, 3(7), 619–627. https://doi.org/10.1016/S2215-0366(16)30065-7

Carhart-Harris, R., & Friston, K. J. (2019). REBUS and the Anarchic Brain: Toward a Unified Model of the Brain Action of Psychedelics. Pharmacological Reviews, 71(3), 316–344. https://doi.org/10.1124/PR.118.017160

Carhart-Harris, R., Giribaldi, B., Watts, R., Baker-Jones, M., Murphy-Beiner, A., Murphy, R., Martell, J., Blemings, A., Erritzoe, D., & Nutt, D. J. (2021). Trial of Psilocybin versus Escitalopram for Depression. New England Journal of Medicine, 384(15), 1402–1411. https://doi.org/10.1056/NEJMOA2032994/SUPPL_FILE/NEJMOA2032994_DATA-SHARING.PDF

Chamberlain, S. R., Fineberg, N. A., Blackwell, A. D., Robbins, T. W., & Sahakian, B. J. (2006). Motor inhibition and cognitive flexibility in obsessive-compulsive disorder and trichotillomania. American Journal of Psychiatry, 163(7). https://doi.org/10.1176/ajp.2006.163.7.1282

Clarke, H. F., Dalley, J. W., Crofts, H. S., Robbins, T. W., & Roberts, A. C. (2004). Cognitive Inflexibility after Prefrontal Serotonin Depletion. Science, 304(5672), 878–880. https://doi.org/10.1126/science.1094987

Clarke, H. F., Walker, S. C., Crofts, H. S., Dalley, J. W., Robbins, T. W., & Roberts, A. C. (2005). Prefrontal serotonin depletion affects reversal learning but not attentional set shifting. The Journal of Neuroscience : The Official Journal of the Society for Neuroscience, 25(2), 532–538. https://doi.org/10.1523/JNEUROSCI.3690-04.2005

Clevenger, S. S., Malhotra, D., Dang, J., Vanle, B., & IsHak, W. W. (2018). The role of selective serotonin reuptake inhibitors in preventing relapse of major depressive disorder. Therapeutic Advances in Psychopharmacology, 8(1). https://doi.org/10.1177/2045125317737264

Daw, N. D. (2009). Trial-by-trial data analysis using computational models.

Del’Guidice, T., Lemay, F., Lemasson, M., Levasseur-Moreau, J., Manta, S., Etievant, A., Escoffier, G., Doré, F. Y., Roman, F. S., & Beaulieu, J. M. (2013). Stimulation of 5-HT2C Receptors Improves Cognitive Deficits Induced by Human Tryptophan Hydroxylase 2 Loss of Function Mutation. Neuropsychopharmacology 2014 39:5, 39(5), 1125–1134. https://doi.org/10.1038/npp.2013.313

Den Ouden, H. E. M., Daw, N. D., Fernandez, G., Elshout, J. A., Rijpkema, M., Hoogman, M., Franke, B., & Cools, R. (2013). Dissociable effects of dopamine and serotonin on reversal learning. Neuron, 80(4), 1090–1100. https://doi.org/10.1016/J.NEURON.2013.08.030

Dias, R., Robbins, T. W., & Roberts, A. C. (1996). Dissociation in prefrontal cortex of affective and attentional shifts. Nature, 380(6569). https://doi.org/10.1038/380069a0

Doss, M. K., Považan, M., Rosenberg, M. D., Sepeda, N. D., Davis, A. K., Finan, P. H., Smith, G. S., Pekar, J. J., Barker, P. B., Griffiths, R. R., & Barrett, F. S. (2021). Psilocybin therapy increases cognitive and neural flexibility in patients with major depressive disorder. Translational Psychiatry, 11(1). https://doi.org/10.1038/s41398-021-01706-y

Fineberg, N. A., Hollander, E., Pallanti, S., Walitza, S., Grünblatt, E., Dell’Osso, B. M., Albert, U., Geller, D. A., Brakoulias, V., Janardhan Reddy, Y. C., Arumugham, S. S., Shavitt, R. G., Drummond, L., Grancini, B., De Carlo, V., Cinosi, E., Chamberlain, S. R., Ioannidis, K., Rodriguez, C. I., … Menchon, J. M. (2020). Clinical advances in obsessive-compulsive disorder: A position statement by the International College of Obsessive-Compulsive Spectrum Disorders. In International Clinical Psychopharmacology. https://doi.org/10.1097/YIC.0000000000000314

Furr, A., Danet Lapiz-Bluhm, M., & Morilak, D. A. (2012). 5-HT2A receptors in the orbitofrontal cortex facilitate reversal learning and contribute to the beneficial cognitive effects of chronic citalopram treatment in rats. International Journal of Neuropsychopharmacology, 15(9). https://doi.org/10.1017/S1461145711001441

Goldberg, S. B., Pace, B. T., Nicholas, C. R., Raison, C. L., & Hutson, P. R. (2020). The experimental effects of psilocybin on symptoms of anxiety and depression: A meta-analysis. Psychiatry Research, 284. https://doi.org/10.1016/J.PSYCHRES.2020.112749

Hervig, M., Fiddian, L., Piilgaard, L., Bozič, T., Blanco-Pozo, M., Knudsen, C., Olesen, S. F., Alsiö, J., & Robbins, T. W. (2020). Dissociable and Paradoxical Roles of Rat Medial and Lateral Orbitofrontal Cortex in Visual Serial Reversal Learning. Cerebral Cortex. https://doi.org/10.1093/cercor/bhz144

Hervig, M., Piilgaard, L., Božic, T., Alsiö, J., & Robbins, T. W. (2020). Glutamatergic and serotonergic modulation of rat medial and lateral orbitofrontal cortex in visual serial reversal learning. Psychology & Neuroscience, 13(3), 438. https://doi.org/10.1037/PNE0000221

Herzallah, M. M., Moustafa, A. A., Natsheh, J. Y., Abdellatif, S. M., Taha, M. B., Tayem, Y. I., Sehwail, M. A., Amleh, I., Petrides, G., Myers, C. E., & Gluck, M. A. (2013). Learning from negative feedback in patients with major depressive disorder is attenuated by SSRI antidepressants. Frontiers in Integrative Neuroscience, 7(SEP). https://doi.org/10.3389/fnint.2013.00067

Iigaya, K., Fonseca, M. S., Murakami, M., Mainen, Z. F., & Dayan, P. (2018). An effect of serotonergic stimulation on learning rates for rewards apparent after long intertrial intervals. Nature Communications 2018 9:1, 9(1), 1–10. https://doi.org/10.1038/s41467-018-04840-2

Izquierdo, A., Carlos, K., Ostrander, S., Rodriguez, D., McCall-Craddolph, A., Yagnik, G., & Zhou, F. (2012). Impaired reward learning and intact motivation after serotonin depletion in rats. Behavioural Brain Research, 233(2), 494–499. https://doi.org/10.1016/J.BBR.2012.05.032

Jentsch, J. D., & Taylor, J. R. (2001). Impaired Inhibition of Conditioned Responses Produced by Subchronic Administration of Phencyclidine to Rats. Neuropsychopharmacology 2000 24:1, 24(1), 66–74. https://doi.org/10.1016/s0893-133x(00)00174-3

Jones, B., & Mishkin, M. (1972). Limbic lesions and the problem of stimulus-Reinforcement associations. Experimental Neurology, 36(2). https://doi.org/10.1016/0014-4886(72)90030-1

Kanen, J. W., Apergis-Schoute, A. M., Yellowlees, R., Arntz, F. E., van der Flier, F. E., Price, A., Cardinal, R. N., Christmas, D. M., Clark, L., Sahakian, B. J., Crockett, M. J., & Robbins, T. W. (2021). Serotonin depletion impairs both Pavlovian and instrumental reversal learning in healthy humans. Molecular Psychiatry 2021 26:12, 26(12), 7200–7210. https://doi.org/10.1038/s41380-021-01240-9

Kanen, J. W., Luo, Q., Kandroodi, M. R., Cardinal, R. N., Robbins, T. W., Nutt, D. J., Carhart-Harris, R. L., & Ouden, H. E. M. den. (2022). Effect of lysergic acid diethylamide (LSD) on reinforcement learning in humans. Psychological Medicine, 1–12. https://doi.org/10.1017/S0033291722002963

Koob, G. F., & Volkow, N. D. (2016). Neurobiology of addiction: a neurocircuitry analysis. The Lancet Psychiatry, 3(8), 760–773. https://doi.org/10.1016/S2215-0366(16)00104-8

Kuypers, K. P. C., Riba, J., de la Fuente Revenga, M., Barker, S., Theunissen, E. L., & Ramaekers, J. G. (2016). Ayahuasca enhances creative divergent thinking while decreasing conventional convergent thinking. Psychopharmacology. https://doi.org/10.1007/s00213-016-4377-8

Liu, F. Y., Xing, G. G., Qu, X. X., Xu, I. S., Han, J. S., & Wan, Y. (2007). Roles of 5-hydroxytryptamine (5-HT) receptor subtypes in the inhibitory effects of 5-HT on C-fiber responses of spinal wide dynamic range neurons in rats. Journal of Pharmacology and Experimental Therapeutics, 321(3). https://doi.org/10.1124/jpet.106.115204

Luo, Q., Kanen, J. W., Bari, A., Skandali, N., Langley, C., Knudsen, G. M., Alsiö, J., Phillips, B. U., Sahakian, B. J., Cardinal, R. N., & Robbins, T. W. (2023). Common roles for serotonin in rats and humans for computations underlying flexible decision-making. BioRxiv, 2023.02.15.527569. https://doi.org/10.1101/2023.02.15.527569

Madsen, M. K., Fisher, P. M., Burmester, D., Dyssegaard, A., Stenbæk, D. S., Kristiansen, S., Johansen, S. S., Lehel, S., Linnet, K., Svarer, C., Erritzoe, D., Ozenne, B., & Knudsen, G. M. (2019). Psychedelic effects of psilocybin correlate with serotonin 2A receptor occupancy and plasma psilocin levels. Neuropsychopharmacology 2019 44:7, 44(7), 1328–1334. https://doi.org/10.1038/s41386-019-0324-9

Marek, G. J., Martin-Ruiz, R., Abo, A., & Artigas, F. (2005). The Selective 5-HT2A Receptor Antagonist M100907 Enhances Antidepressant-Like Behavioral Effects of the SSRI Fluoxetine. Neuropsychopharmacology 2005 30:12, 30(12), 2205–2215. https://doi.org/10.1038/sj.npp.1300762

Michely, J., Eldar, E., Erdman, A., Martin, I. M., & Dolan, R. J. (2022). Serotonin modulates asymmetric learning from reward and punishment in healthy human volunteers. Communications Biology, 5(1). https://doi.org/10.1038/s42003-022-03690-5

Murphy-Beiner, A., & Soar, K. (2020). Ayahuasca’s ‘afterglow’: improved mindfulness and cognitive flexibility in ayahuasca drinkers. Psychopharmacology. https://doi.org/10.1007/s00213-019-05445-3

Perani, D., Garibotto, V., Gorini, A., Moresco, R. M., Henin, M., Panzacchi, A., Matarrese, M., Carpinelli, A., Bellodi, L., & Fazio, F. (2008). In vivo PET study of 5HT(2A) serotonin and D(2) dopamine dysfunction in drug-naive obsessive-compulsive disorder. NeuroImage, 42(1), 306–314. https://doi.org/10.1016/J.NEUROIMAGE.2008.04.233

Phillips, B. U., Dewan, S., Nilsson, S. R. O., Robbins, T. W., Heath, C. J., Saksida, L. M., Bussey, T. J., & Alsiö, J. (2018). Selective effects of 5-HT2C receptor modulation on performance of a novel valence-probe visual discrimination task and probabilistic reversal learning in mice. Psychopharmacology. https://doi.org/10.1007/s00213-018-4907-7

Pokorny, T., Duerler, P., Seifritz, E., Vollenweider, F. X., & Preller, K. H. (2020). LSD acutely impairs working memory, executive functions, and cognitive flexibility, but not risk-based decision-making. Psychological Medicine, 50(13), 2255–2264. https://doi.org/10.1017/S0033291719002393

R Core Team. (2021). R: A Language and Environment for Statistical Computing. In R Foundation for Statistical Computing.

Roth, B. L. (2011). Irving Page Lecture: 5-HT(2A) serotonin receptor biology: interacting proteins, kinases and paradoxical regulation. Neuropharmacology, 61(3), 348–354. https://doi.org/10.1016/J.NEUROPHARM.2011.01.012

Rygula, R., Clarke, H. F., Cardinal, R. N., Cockcroft, G. J., Xia, J., Dalley, J. W., Robbins, T. W., & Roberts, A. C. (2015). Role of central serotonin in anticipation of rewarding and punishing outcomes: Effects of selective amygdala or orbitofrontal 5-HT Depletion. Cerebral Cortex, 25(9), 3064–3076. https://doi.org/10.1093/cercor/bhu102

Santana, N., Bortolozzi, A., Serrats, J., Mengod, G., & Artigas, F. (2004). Expression of serotonin1A and serotonin2A receptors in pyramidal and GABAergic neurons of the rat prefrontal cortex. Cerebral Cortex, 14(10). https://doi.org/10.1093/cercor/bhh070

Seymour, B., Daw, N. D., Roiser, J. P., Dayan, P., & Dolan, R. (2012). Serotonin selectively modulates reward value in human decision-making. Journal of Neuroscience, 32(17). https://doi.org/10.1523/jneurosci.0053-12.2012

Stroud, J. B., Freeman, T. P., Leech, R., Hindocha, C., Lawn, W., Nutt, D. J., Curran, H. V., & Carhart-Harris, R. (2018). Psilocybin with psychological support improves emotional face recognition in treatment-resistant depression. Psychopharmacology, 235(2), 459. https://doi.org/10.1007/S00213-017-4754-Y

Sutton, R. S., & Barto, A. G. (1998). Reinforcement Learning: An Introduction. IEEE Transactions on Neural Networks, 9(5), 1054–1054. https://doi.org/10.1109/tnn.1998.712192

Torrado Pacheco, A., Olson, R. J., Garza, G., & Moghaddam, B. (2023). Acute psilocybin enhances cognitive flexibility in rats. Neuropsychopharmacology. https://doi.org/10.1038/s41386-023-01545-z

Uddin, L. Q. (2021). Cognitive and behavioural flexibility: neural mechanisms and clinical considerations. In Nature Reviews Neuroscience (Vol. 22, Issue 3, pp. 167–179). Nature Research. https://doi.org/10.1038/s41583-021-00428-w

Wickham, H. (2014). Tidy Data. Journal of Statistical Software, 59(10), 1–23. https://doi.org/10.18637/JSS.V059.I10

Zhu, C., Kwok, N. T. kit, Chan, T. C. wan, Chan, G. H. kei, & So, S. H. wai. (2021). Inflexibility in Reasoning: Comparisons of Cognitive Flexibility, Explanatory Flexibility, and Belief Flexibility Between Schizophrenia and Major Depressive Disorder. Frontiers in Psychiatry, 11, 609569. https://doi.org/10.3389/FPSYT.2020.609569/BIBTEX

Zühlsdorff, K., López-Cruz, L., Dutcher, E. G., Jones, J. A., Pama, C., Sawiak, S., Khan, S., Milton, A. L., Robbins, T. W., Bullmore, E. T., & Dalley, J. W. (2023). Sex-dependent effects of early life stress on reinforcement learning and limbic cortico-striatal functional connectivity. Neurobiology of Stress, 22, 100507. https://doi.org/10.1016/J.YNSTR.2022.100507

