## Supplementary materials and results for "5-HT 2A and 5-HT 2C receptor antagonism differentially modulates reinforcement learning and cognitive flexibility: behavioral and computational evidence"

**Methods**

**Drugs**

M100907 (R-(+)-α-(2,3-dimethoxyphenyl)-1-[2-(4-fluorophenylethyl)]-4-piperidinemethanol) (Sigma Aldrich, #M3324), a highly selective 5-HT_2A_R antagonist (Kehne et al., 1996), was dissolved in 0.01 M phosphate-buffered saline (PBS) and 0.1 M hydrochloride, and adjusted with NaOH to pH 7. M100907 was administered at 0 (vehicle), 0.03 or 0.1 mg/kg.

SB-242084 (Eli Lilly, Indianapolis, IN, USA) was first dissolved in polyethene glycol 400 (PEG400) (Fisher Scientific, Loughborough, UK) at 20% of the final required volume, and then made up by 10% (w/v) hydroxypropyl-beta-cyclodextrin (Sigma-Aldrich, Poole, UK) in saline, and checked that the pH was 7. For systemic treatment, SB-242084 was administered intraperitoneally (i.p.) at doses of 0 (vehicle), 0.3 or 1.0 mg/kg in a volume of 1 ml/kg, 30 min prior to testing. Drugs were divided into the aliquots required for each test day and frozen at −80°C.

**Apparatus**

Animals were trained and tested in 12 operant chambers with touchscreens (Med Associates, Georgia, VT, USA) placed in sound- and light-attenuating wooden cabinets equipped with a fan for ventilation and masking of external noise. The chambers measured 30×39×29 cm and consisted of a clear Perspex ceiling, front door and back panel, and metal paneling on the sides of the chamber. A metal grid with a removable metal tray below made up the floor of the chamber. A central food magazine coupled to an external pellet dispenser was located on one side of the chamber. It was equipped with a light and infrared beam sensors to detect magazine entry and allowed delivery of one 45 mg sucrose pellet (TestDiet 5TUL; Sandown Scientific, Middlesex, UK) upon correct responses. A house-light (~3 W) was located near the ceiling directly above the magazine. A touch-sensitive screen (29×32 cm; (Elo Touch Solutions, Inc.) presenting visual stimuli was located on the side opposite to the magazine. Chambers were controlled by in-house software (Visual Basic 2010 Express.NET, Microsoft 2010, (Mar et al., 2013)) and running task schedules previously published (Alsiö et al., 2019).

**Behavioral pre-training**

The initial five-stage pre-training phase has been described in detail previously (Hervig et al., 2020). Following food restriction, pre-training began involving Pavlovian and instrumental conditioning. This was followed by visual discrimination training with two visual stimuli, as well as visual discrimination and reversal learning including an additional third (probe) stimulus. In stages 1 to 3, rats were trained to respond to a single white square stimulus at the bottom center of the touchscreen for sucrose reward pellets, during 60-minute daily sessions, until they reached a criterion of receiving the maximum 100 pellets in one session. The stimulus decreased in size across the three stages until a final size of 3×4 cm (‘start box’) in stage 3. In pre-training stages 4 and 5, two additional stimuli were introduced (horizontal and vertical bars); first at the bottom of the screen to ease touch (stage 4), then the stimulus was raised 5 cm to the final location on the screen to avoid accidental touches (stage 5). At this stage, touching the white starting stimulus was no longer reinforced, but instead led to the presentation of one of these novel stimuli to the left or right (pseudo-randomized location). Responding to the presented stimulus was reinforced with a sugar pellet, whereas responding to the blank side was signaled as incorrect by the illumination of the house-light for a 5 s time-out period. Performance at ≥80% correct touches to one stimulus in a session led to training sessions with the other stimulus. When criterion of >80% correct touches was reached also on this stimulus, the rat moved on from stage 4 to stage 5, and after ≥80% correct touches were reached on both stimuli on stage 5, visual discrimination training ensued.

**Touchscreen visual discrimination and reversal**

During the visual discrimination stage, rats were presented with both stimuli simultaneously, of which one was reinforced. For session initiation, the rats would collect a free reward delivery, which led to presentation of the start box. The rat initiated a trial by responding to the start box, which initiated a simultaneous presentation of the stimuli pair (CS+ vs. CS−; ‘horizontal bars’ or ‘vertical bars’; counterbalanced across rats). Responding to the correct stimulus (A+) was reinforced with a sugar pellet, while responding to the incorrect non-reinforced stimulus (B–) triggered a house-light-signaled 5 s time-out period. Failure to make a choice of either stimulus within 10 s of trial initiation was recorded as an omission. A 5 s inter-trial-interval (ITI) period preceded the next trial. To prevent the rats from developing a side bias, the stimuli were presented on the screen (left or right side) in a pseudo-random order (max. 3 consecutive trials to the same side). The daily session ended after either 60 min, 150 rewards, or 250 trials, whichever was the first to occur. The rats reached criterion performance by making 24 correct responses out of a running window of 30 trials. Prior to the reversal learning training, a retention session with the same reward contingencies was given, as well as on the day following attainment of the learning criterion, to ensure that the rat had acquired the discrimination.

Following the retention session during visual discrimination, the contingencies reversed so the rats now had to respond to the previous non-rewarded CS− stimulus (now CS+) for reinforcement until they reached the reversal learning criterion (24/30). A retention session both preceded and followed a reversal block.

**Hierarchical Bayesian reinforcement learning modelling**

In the highest hierarchical level, there was a group-specific mean and standard deviation for each of the parameters. **Supplementary** **Table 1** provides a summary of the prior values for these parameters. The parameters were drawn for each subject from a normal distribution with the respective mean and standard deviation. An RL algorithm was fitted to the behavior, and the highest posterior density interval (HDI) was calculated for group mean differences (Kruschke, 2014). Please see the main text for an explanation of how the Q-values were updated and associated equations.

The models that were fitted to the data were the following:

1. **Two parameters:** **α and β**, the learning rate and reinforcement sensitivity parameter. The Q-value on each trial is calculated by adding the Q-value for that stimulus from the previous trial to the prediction error multiplied by the learning rate. See equation 1 in the main text for further information.

2. **Three parameters: α,** **β,** stimulus stickiness parameter **κ_stim_**. The κ_stim_ parameter represents the tendency to respond to the same stimulus as on the previous trial, regardless of its location and whether the stimulus was rewarded or not. This parameter updates the Q-value according to: ${Q^{stim}}_{s,t}=\kappa_{stim}S_{s,t-1}$, where $S_{s,t-1}$ is the stimulus chosen by the subject on the previous trial. This value is 1 if the same stimulus was chosen, and 0 if the other stimulus was chosen. Finally, the Q-value is determined by the sum of ${Q^{stim}}_{s,t}$ and the Q-value as calculated in equation 1.

3. **Three parameters: α,** **β,** side stickiness parameter **κ_side_**. Instead of the stimulus stickiness parameter, the side stickiness parameter was included. It reflects the tendency to select the same side as on the last trial. It was used to update the Q-values using the following equation: ${Q^{loc}}_{s,t}=\kappa_{side}L_{s,t-1}$. $L_{s,t-1}$ is the side chosen by the subject on the last trial. This value is 1 if the same side was chosen, and 0 if the other side was chosen. The final Q-value is the sum of ${Q^{loc}}_{s,t}$ and the Q-value as calculated in equation 1.

4. **Four parameters: α, β, κ_stim_ and κ_side_.**

5. **Three parameters: α_rew_, α_pun_, β.** As in model 1, but the learning rates were separated for rewarded and non-rewarded trials.

6. **Four parameters: α_rew_, α_pun_, β and κ_stim_.**

7. **Four parameters: α_rew_, α_pun_, β and κ_side_.**

8. **Five parameters: α_rew_, α_pun_, β, κ_stim_ and κ_side_.**

9. **Six parameters: α_rew_, α_pun_, β, κ_stim_, κ_side_, discount factor ρ**. The novel parameter **ρ** was included to better reflect behavior on the VPVD task. It accounts for slower learning from the probe stimulus, as this stimulus is randomly linked to reward. Note also that the task was designed such that the rats had extensive experience with the probe stimulus during pre-training. The value updating is only discounted if the probe stimulus has been chosen. During a trial on which the probe stimulus is presented, the value change is calculated according to: $\left( \alpha\times\left( r_{t}-Q_{t}\left( c_{t} \right) \right) \right)/\left( 1+\rho\right)$. The prior distribution of ρ was gamma(α=4.82, β=0.88) and it had a lower bound of 0. The learning rate in this case can either be $\alpha_{rew}$ or $\alpha_{pun}$, depending on the feedback received.

Hamiltonian Markov chain Monte Carlo sampling was used to fit the models via Stan 2.17.2 (Carpenter et al., 2017). The potential scale reduction factor was used to ensure convergence (Brooks & Gelman, 1998; Gelman, 2013). A value close to 1 indicates perfect convergence. A cut-off of 1.1 was selected as a stringent criterion for convergence. Models were compared using a bridge sampling estimate of the marginal likelihood using the “bridgesampling” R package (Gronau et al., 2017, 2020).

Given that the simulations for the 5-HT_2C_R receptor antagonist (SB-242084, Experiment 2) data from model 7 did not fully capture the animals’ original behaviour on the task (see **Supplementary Figure 2** and **Fig. 3A**), and that model 9 best explained the 5-HT_2A_R antagonist (M100907, Experiment 1) data, model 9 performance was examined in more detail, given that this was the winning model for Experiment 1 but did not converge well for this data set. First, the Stan parameters *adapt_delta* and *max_treedepth* were increased and the step size was decreased when running this model in an attempt to attain convergence; however, this did not result in convergence. Next, animals that did not reach criterion of 80% were excluded (5 rats). Nonetheless, convergence was not achieved, perhaps because performance of the excluded rats was still high (at above 70%). Lastly, the prior for the $\rho$ parameter was changed to a beta(1.2, 1.2) distribution, to tighten the prior. This model did also not converge. Therefore, we proceeded with the winning model, model 7.

To check that the results from model 7 would be the same as for model 9 for the 5-HT_2A_R antagonist data, group comparisons were also analyzed for this dataset. Indeed, it confirmed that the results would be the same; the punishment learning rate was decreased at the lower dose of M100907 (group difference, 0 ∉ 95% HDI), as well as at the higher dose (group difference, 0 ∉ 75% HDI). The reinforcement sensitivity was also decreased at the lower dose (group difference, 0 ∉ 75% HDI) and the side stickiness was increased at this dose (group difference, 0 ∉ 95% HDI).

This was also repeated for the 5-HT_2C_R antagonist dataset (i.e., the group comparisons of model 9 were analyzed despite it not converging), and the results were also the same. At the low dose of SB-242084, reinforcement sensitivity was lower compared to the control group (group difference, 0 ∉ 75% HDI). Side stickiness was reduced at both low and high doses (group difference, 0 ∉ 95% HDI and group difference, 0 ∉ 75% HDI, respectively). No differences were found in the other parameters.

**Results**

**Additional analyses of conventional measures**

There was a trend for M100907 to decrease win-stay probability without reaching statistical significance (*F*_2,35_ = 2.91, *p* = 0.068) with planned pairwise comparison revealing 0.03 mg/kg M10907 marginally reducing win-stay probability (*t*_38.3_ = -2.30, *p* = 0.051) (**Figure SF.3)**. Likewise, M100907 trended towards decreasing lose-shift probability (*F*_2,35_ = 3.04, *p* = 0.061) with a planned pairwise comparison for 0.03 mg/kg M10907 indicating significantly reduced lose-shift probability (*t*_38.3_ = -2.32, *p* = 0.048).

We found no effects of M100907 on response or collection latencies.

SB-242084 affected win-stay probability significantly (*F*_2,35_ = 3.50, *p* = 0.041) with planned pairwise comparison revealing that there was a trend for 1 mg/kg SB-242084 to reduce win-stay probability, although without reaching significance (*t*_38.3_ = -2.29, *p* = 0.053) (**Figure SF.4)**. Likewise, SB-242084 affected lose-shift probability significantly (*F*_2,35_ = 3.36, *p* = 0.046) with a planned pairwise comparison revealing that there was a trend for 1 mg/kg SB-242084 to reduce win-stay probability, albeit non-significantly (*t*_38.3_ = -2.11, *p* = 0.078).

SB-242084 did not affect response latencies but there was a trend towards speeding collection latencies (*F*_2,35_ = 2.69, *p* = 0.082).

**Supplementary Table 1.** Priors for model parameters.

| **Parameter** | **Prior** | **Reference** |
| --- | --- | --- |
| Reward learning rate, $\alpha_{rew}$ | Beta(1.2, 1.2) | (Den Ouden et al., 2013) |
| Punishment learning rate, $\alpha_{pun}$ | Beta(1.2, 1.2) | (Den Ouden et al., 2013) |
| Combined learning rate, $\alpha$ | Beta(1.2, 1.2) | (Den Ouden et al., 2013) |
| Reinforcement sensitivity, β | Gamma(α=4.82, β=0.88) | (Gershman, 2016) |
| Side stickiness, κside | Normal(0,1) | (Christakou et al., 2013) |
| Stimulus stickiness, κstim | Normal(0,1) | (Christakou et al., 2013) |
| Discount factor, ρ | Gamma(α=4.82, β=0.88) | (De novo) |
| **Intersubject variability in parameters** |  |  |
| $\alpha_{rew}$, $\alpha_{pun}$, $\alpha$, κside, κstim intersubject standard deviations | Normal(0,0.05) constrained to $\geq$0 | (Kanen et al., 2019) |
| β intersubject standard deviations | Normal(0,1) constrained to $\geq$0 | (Gershman, 2016) |

**Supplementary Table 2.** Mean and standard deviation of ρ values from model 9 fitted to the M100907 dataset.

| Group | Mean | Standard deviation |
| --- | --- | --- |
| Control | 4.82 | 1.84x10^-2^ |
| Low dose | 8.49 | 4.85x10^-2^ |
| High dose | 5.00 | 2.19x10^-2^ |

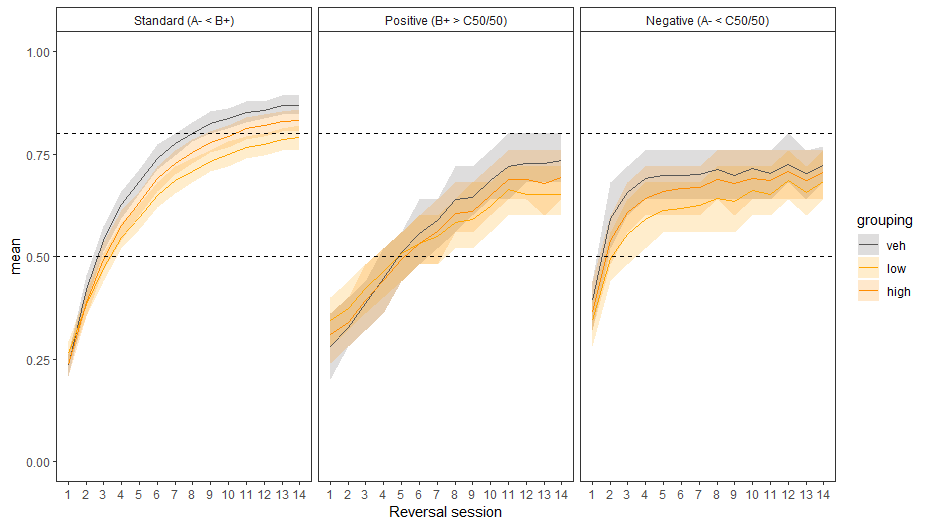
**Supplementary Figure 1.** Behavioral simulations for the M100907 dataset using the winning model parameters (vd, standard visual discrimination trials; pos, positive probe trials; neg, negative probe trials; veh, vehicle). The ordinate (y axis) represents the mean for that behavioral measure across all simulated subjects. See **Figure 2A** for a direct comparison to measures from the VPVD task.

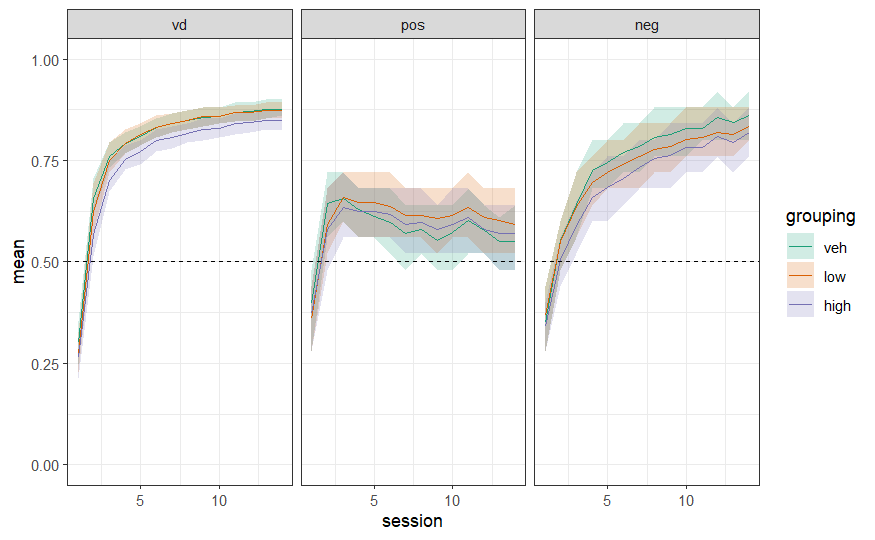
**Supplementary Figure 2.** Behavioral simulations for the SB-242084 dataset using the winning model parameters (vd, standard visual discrimination trials; pos, positive probe trials; neg, negative probe trials; veh, vehicle). The ordinate (y axis) represents the mean for that behavioral measure across all simulated subjects. See **Figure 3A** for direct comparison to measures from the VPVD task.

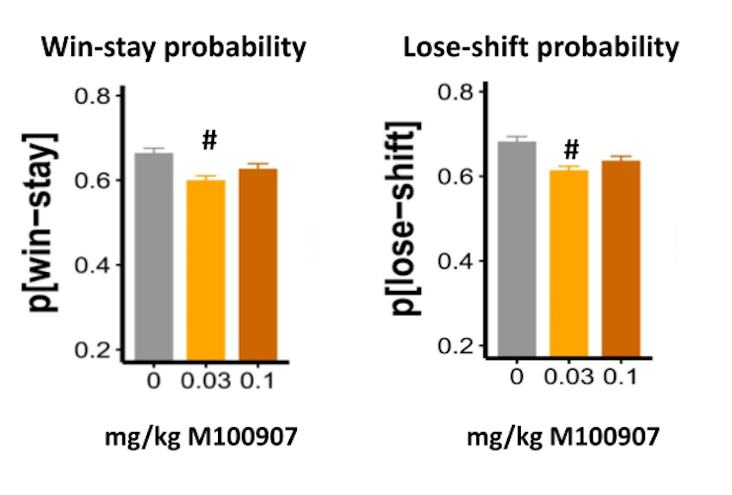
**Supplementary Figure 3.** Effects of M100907 on the conventional VPVD measures win-stay and lose-shift probabilities. Results are represented as mean ± standard error of the mean (SEM); *** *p* < 0.01, # p < 0.1.

**Supplementary Figure 4.** Effects of SB-242048 on the conventional VPVD measures win-stay and lose-shift probabilities. Results are represented as mean ± standard error of the mean (SEM); *** *p* < 0.01, # p < 0.1.

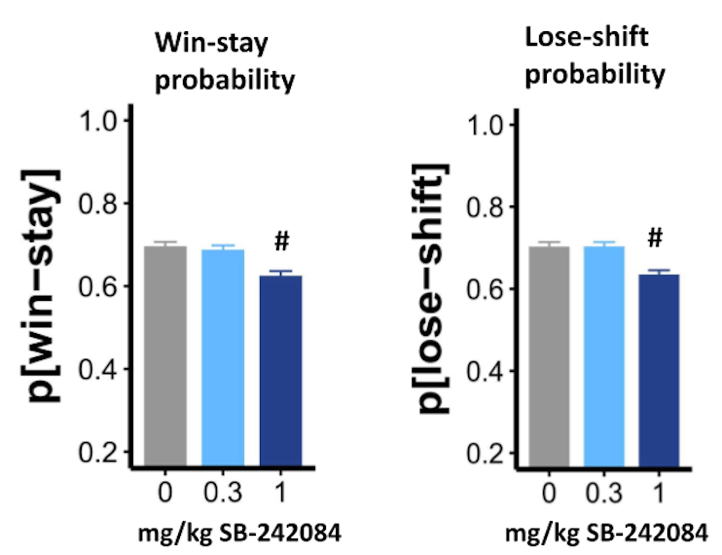
